# Substrate Resistance to Traction Forces Controls Fibroblast Polarization

**DOI:** 10.1101/2020.05.15.098046

**Authors:** D. Missirlis, T. Haraszti, L. Heckmann, J. P. Spatz

## Abstract

The mechanics of fibronectin-rich extracellular matrix regulate cell physiology in a number of diseases, prompting efforts to elucidate cell mechanosensing mechanisms at the molecular and cellular scale. Here, the use of fibronectin-functionalized silicone elastomers that exhibit considerable frequency-dependence in viscoelastic properties unveiled the presence of two cellular processes that respond discreetly to substrate mechanical properties. Soft elastomers supported efficient focal adhesion maturation and fibroblast spreading due to an apparent stiff surface layer. However, soft elastomers did not enable cytoskeletal and fibroblast polarization; elastomers with high cross-linking and low deformability were required for polarization. The underlying reason for this behavior was the inability of soft elastomeric substrates to resist traction forces, rather than a lack of sufficient traction force generation; accordingly, mild inhibition of actomyosin contractility rescued fibroblast polarization even on the softer elastomers. Our findings help reconcile previously proposed local and global models of cell mechanosensing by demonstrating the differential dependence of substrate mechanics on distinct cellular processes.

**Statement of Significance:** The mechanisms cells employ to sense and respond to the mechanical properties of their surroundings remain incompletely understood. In this study we used a commercial silicone elastomer formulation to prepare compliant, fibronectin-coated substrates and investigate the adhesion and polarization of human fibroblasts. Our results suggest the existence of at least two discrete mechanosensing processes regulated at different time and length (force) scales. Focal adhesion assembly and cell spreading were promoted by a stiff surface layer independent from bulk viscoelasticity, whereas effective cell polarization required elevated elastomer stiffness, sufficient to resist applied cell traction. The results presented here have implications on the use of elastomeric substrates as biomaterials for mechanosensing studies or clinical applications.

## Introduction

Cells engage their microenvironment by using specialized receptors that recognize biochemical ligands presented on the extracellular matrix (ECM). The result of ligand-receptor interactions depends on the physical connection of these ligands to their environment: the outcome can differ whether the ligand is soluble compared to being attached to another cell or the ECM, and thus the ligand-receptor pair being subject to force (1). The field of mechanosensing has emerged to characterize the processes that enable cells to interpret physical forces that are applied on the cell, or that result from resistance of cell-generated forces on the external environment (2, 3). The interplay of biochemical and physical stimuli during mechanosensing is invariably intertwined: how cells sense their surrounding mechanical environment is a function of which ligands are presented and which receptors bind them.

Over the past two decades, numerous demonstrations that alteration of surrounding mechanics is associated with fundamental cell processes (4), fate (5) and pathologies (6, 7), have prompted scientists to adopt reductionist, in vitro experimentation in order to fundamentally understand mechanosensing (8). Engineered hydrogels and silicone elastomers are popular substrate choices, as they are easy to fabricate, provide straightforward control over viscoelastic properties and enable direct, high-resolution observations. However, while the majority of studies have validated the view that substrate elasticity controls cell behavior, important differences were described between materials of similar reported mechanical properties (9-12).

There are *four* major reasons for the observed differences on cell behavior when cells are interrogated on substrates of equal nominal stiffness but dissimilar material composition. *First*, different mechanical characterization techniques probe distinct properties and at different scales. Thus, resulting stiffness values can vary for the same material (13) and the use of a single nominal stiffness to characterize the substrate can confound conclusions.

*Second*, mechanosensing depends on the biochemical properties of the presented ligands as well as the cell receptors and intracellular composition, rendering mechanosensing sensitive to cell and biochemical factors (14-16). *Third*, ligand tethering mechanics can modulate force loading rates, and therefore associated mechano-transducing events (17). For example, if the link between ligand and substrate cannot sustain sufficient force, cells are unable to adhere independent of the stiffness of the underlying substrate (18). Besides this extreme, binary case, a range of potential force-bearing linkages can exist, which have been shown to affect force loading (19-21). Moreover, traction force-induced changes in conformation of large ECM proteins such as fibronectin (FN) can alter receptor-ligand engagement and dynamics (22). *Fourth*, the often-neglected viscous/plastic component of the substrate can modulate force loading rates and affect ligand density as cells pull towards them new ligands and form extra bonds (23-28).

Given the above considerations, independent control over cell ligand coupling to the underlying substrate and viscoelastic properties of that substrate is desirable to uncouple the relative contribution of both parameters. Hydrogels typically present covalently-linked ligands to cells, since adsorption of ECM proteins or adhesive peptides onto hydrophilic interfaces is unfavorable. While there is progress in gaining better control over immobilization chemistries (29, 30), there are only few reports on non-covalent binding of ligands on hydrogels (31-33). On the other hand, the hydrophobic nature of silicone elastomers allows for efficient adsorption of ECM proteins and their surface can be modified to additionally enable covalent coupling, albeit with the risk of concurrent modifications in the physicochemical properties of its upper layer (34, 35).

In this study, we investigated the interactions of primary fibroblasts with fibronectin-coated substrates, which were produced with the silicone elastomer formulation CY52-276. Even though this formulation has been employed for mechanosensing studies due to a reported Young’s modulus similar to that of tissues at the lower kPa range (36-39), a comprehensive characterization of its viscoelastic properties is lacking and variation of the mechanical properties through modulation of cross-linking was not explored. In particular, previous studies have relied on characterization of bulk viscoelastic properties, often reporting a single Young’s modulus value to characterize the substrate. Here, we modulated the ratio between base polymer and curing agent, producing substrates that exhibited frequency-dependent viscoelasticity, spanning a physiologically-relevant range of mechanical properties. Furthermore, differences in surface mechanics were explored, revealing the critical role of solid surface tension and its effects on cell adhesion. The possibility of non-covalent fibronectin functionalization of silicone elastomers allowed us to address how potential ECM protein remodeling in conjunction with substrate viscoelasticity affect cell adhesion and polarization. The focus was placed on fibronectin due to the paramount importance of its presence and remodeling in guiding cell behavior during physiological and pathological processes in vivo (40-42). Our findings shed light on the mechanisms that cells utilize to probe substrate mechanics and indicate that there are two relevant length and time scales that govern focal adhesion (FA) maturation and cell polarization.

## Results

### Bulk viscoelastic properties of ultra-soft elastomers

The viscoelastic properties of the silicone elastomer formulation CY52-276 were controlled by the weight ratio (ζ) between the base (component A) and the cross-linker (component B). Lower and higher ratios, compared to the recommended ζ=B/A=1.0, were prepared to obtain “softer” and “stiffer” substrates, respectively. Reducing ζ below 0.7, resulted in a predominantly viscous, tacky and difficult to handle material; therefore, the elastomer with ζ=0.7 was the softest formulation examined. Modulation of mechanical properties and the elastic character of the elastomers were evident by visual inspection after poking with a pipette tip (**Supplementary Movie 1**). Silicone elastomers formed readily on multiple substrates, including glass or tissue culture polystyrene (TCPS), and their thickness could be controlled from tens to hundreds of micrometers by regulating the rotational speed during spin coating (**Supplementary Fig. S1A**). Elastomers thicker than 100 μm were used for all cell experiments.

Oscillatory rheology was used to monitor bulk elastomer properties (**Fig. 1**). The two elastomer components were thoroughly mixed and degassed to remove entrapped air, before application between two parallel plates of the rheometer. The temperature was then set at 65°C and the cross-linking kinetics were monitored over time at a frequency of 1 Hz and 5% strain. The shear storage and loss moduli increased rapidly the first few minutes and reached a plateau within the first 3 hours (**Supplementary Fig. S1B**). All subsequent samples were prepared by heating elastomers at 65°C in an oven for 3 hours.

**Figure 1.**
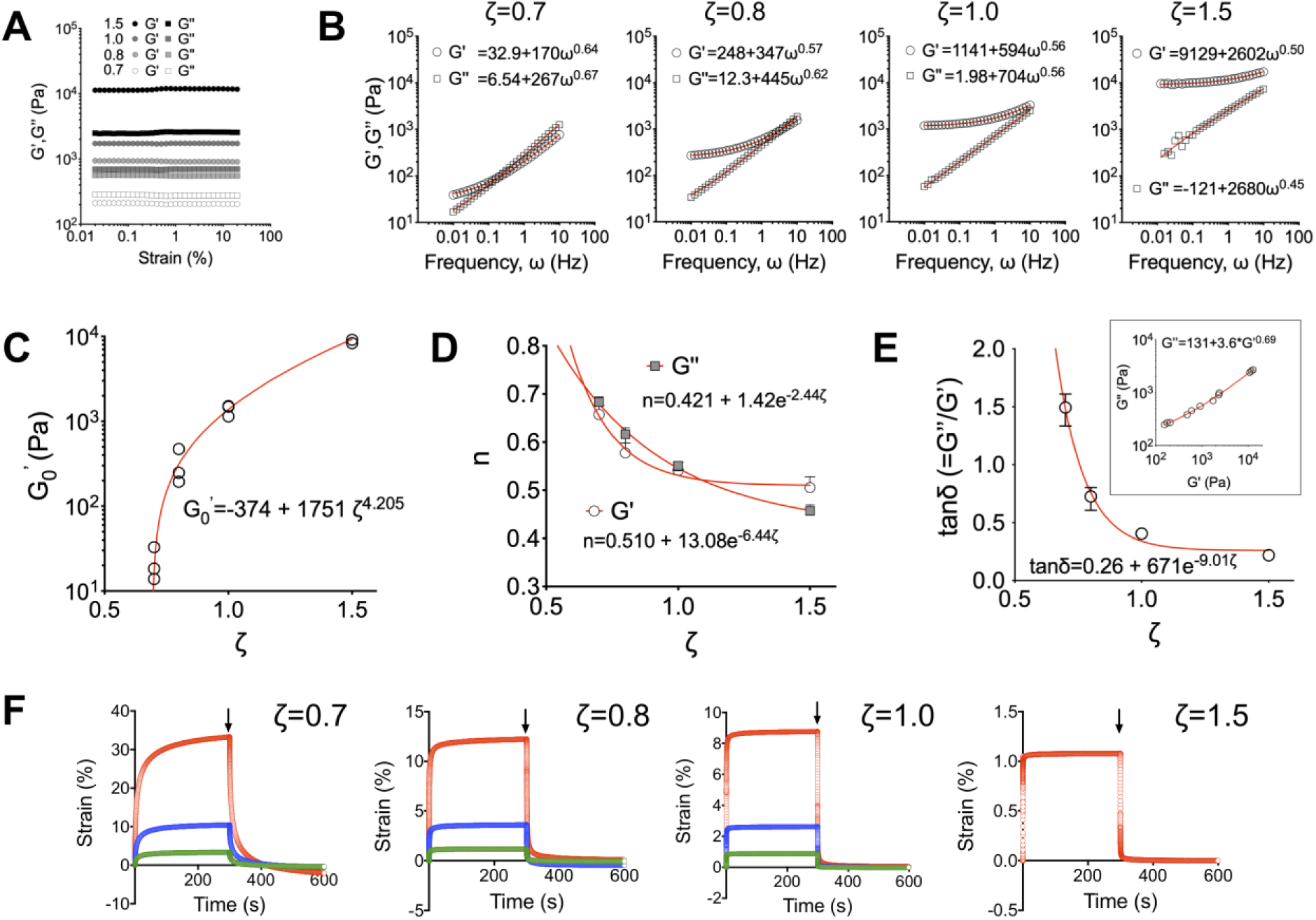
Control over bulk viscoelasticity of silicone elastomers through the ratio of base polymer to cross-linker. **A)** Storage (*G*^*′*^) and loss (*G*^*′′*^) moduli measured at a frequency of 1 Hz are independent of strain in the linear viscoelastic regime. **B)** *G*^*′*^and *G*^*′′*^ as a function of frequency for different ζ ratios. **C)** Extrapolated values of storage modulus for zero frequency 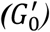 as a function of ζ. Data from N=3 independent experiments are shown. **D)** Dependence of exponent n (from *G*^*′*^, *G*^*′′*^ = a + bω^*n*^) on ratio ζ. Average values and range from N=3. **E)** Dependence of dissipation factor on ratio ζ (Average values and range from N=3); inset shows the relationship between *G*^*′*^and *G*^*′′*^ for data obtained from all elastomers and ζ ratios. *G*^*′*^and *G*^*′′*^ presented were obtained from measurements at 1 Hz frequency and 1% strain. **F)** Creep experiments for elastomers with different ζ values and different imposed stresses. Following stress removal, the elastomers relax to their original positions, indicating lack of bulk plastic deformations.

Elastomers exhibited a linear viscoelastic regime in the range of 0.1 to 10% strain (**Fig. 1A**). The frequency-dependent storage (*G*^*′*^) and loss (*G*^*′′*^) moduli followed a power-law behavior (*G*^*′*^, *G*^*′′*^∼ω^*n*^) with the storage modulus approaching a plateau at the low frequency limit (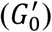), typical for cross-linked elastomers (43) (**Fig. 1B**). Plotting 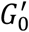 as a function of ζ revealed the dependence of equilibrium elasticity as a function of ζ, confirming that ζ=0.7 is the lower limit for producing viscoelastic solids (**Fig. 1C**). The obtained value of 3.4 kPa for ζ=1 is very close to the one previously reported (44). The exponent n for both *G*^*′*^ and *G*^*′′*^ decreased with ζ, reflecting the changes in cross-linking density (43) (**Fig. 1D**). Importantly, the loss modulus was comparable to the storage modulus for low ζ, with the dissipation factor tan*d=G*^*′′*^*/G*^*′*^ reaching values >1 at 1% strain and *ω*=1 Hz (**Fig. 1E**). The high viscous component of the elastomers raised concerns over potential plastic deformations of the material. However, despite a significant creep behavior for the softer elastomers, there was no evidence of plastic elastomer deformation for strains up to 30% (**Fig. 1F)**. These data indicate complete connectivity of the polymer network at the macroscale.

Silicone elastomers are solvent-free, but may contain an important number of non-crosslinked polymer chains or low molecular weight oligomers, especially as the ratio of cross-linker (ζ) decreases. Indeed, the sol fraction of elastomers increased from 0.37 for ζ=1.5, to 0.52 for ζ=1.0 to 0.67 for ζ=0.8 to 0.71 for ζ=0.7 (mean of two independent experiments; n=3/experiment).

Overall, control over elastomer mechanical properties was feasible through adjustment of the ratio of cross-linker to base polymer, ζ. Despite their important viscous component and high sol fractions, soft elastomers did not exhibit bulk viscoplasticity. Given the large frequency dependence in moduli, we have opted to avoid the use of nominal stiffness values for the rest of the manuscript; instead, we will express our results as a function of ζ, referring to elastomers with low (0.7-0.8) or high (1.0-1.5) ζ as “soft” and “stiff” substrates, respectively.

### Solid surface tension contributes to surface mechanical properties

The surface mechanical properties of silicone elastomers were characterized by an additional technique: indentation measurements using micron-sized colloidal probes and atomic force microscopy (AFM). Initial attempts using spherical glass indenters, even when coated with inert bovine serum albumin (BSA), on untreated elastomers were unsuccessful due to the very high adhesion forces observed between tip and elastomer (44, 45). Measurements performed in ethanol revealed large adhesion forces and a jump-to-contact, rendering analysis using standard models problematic (**Supplementary Fig S2A**). Considering that the targeted application of these materials as cell substrates entails their coating with cell adhesive ECM proteins, we opted to perform the AFM characterization on BSA-coated elastomers, which effectively eliminated non-specific adhesion and thus allowed for AFM indentation measurements. Of note, we avoided plasma-or UV-treatment to render the elastomer surface hydrophilic through oxidation, since such treatments can result in the creation of thin, brittle oxide films exhibiting much higher stiffness compared to the bulk (46, 47).

Force-distance (F-d) curves from indentation on BSA-coated elastomers were initially fitted using the classical Hertz model. The apparent Young’s modulus (*E*) of elastomers increased with ζ, for a fixed indentation force of 10 nN and an indentation speed of 1 μm/s, as expected (**Fig. 2A**). Consistent with the viscoelastic character of the elastomers (**Fig. 1**), a decrease in indentation speed resulted in lower Young’s moduli that approached a plateau value (**Fig. 2B**). Accordingly, hysteresis was evident, even at an indentation speed of 0.1 μm/s, which was the lowest studied as a compromise between experimental time and simulation of equilibrium conditions (**Supplementary Fig. S2B,C**). Analysis of the F-d curves up to different indentation forces resulted in an apparent Young’s modulus as a function of indentation depth. Interestingly, the calculated modulus increased with decreasing indentation for low values of ζ, while the dependence was lost, and in some experiments inversed, at higher ζ values (**Fig. 2C**).

**Figure 2.**
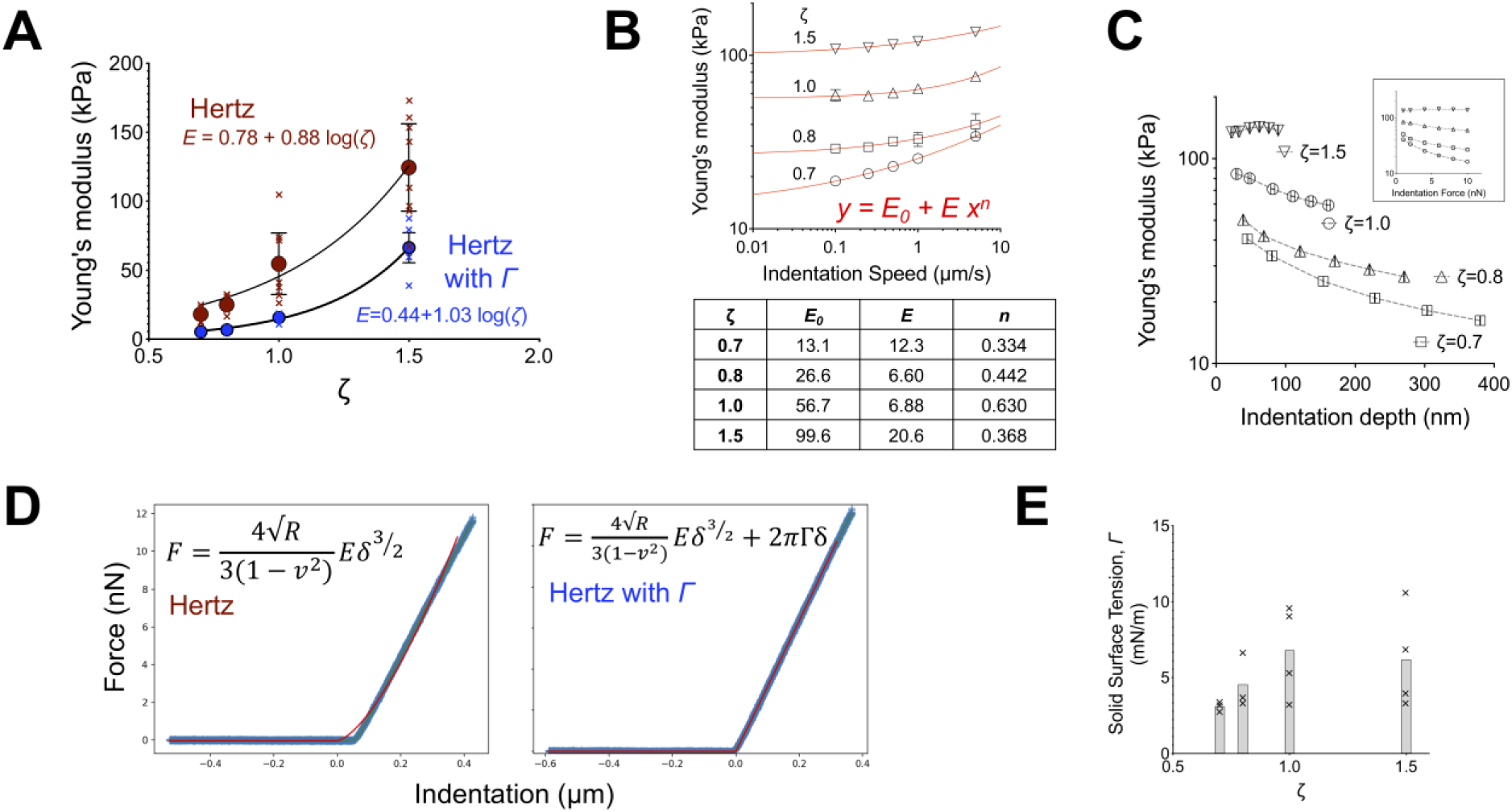
The surface layer of silicone elastomers exhibits higher apparent stiffer due to solid surface tension. **A)** Apparent Young’s moduli determined by fitting F-d curves from AFM indentation measurements with the standard Hertz model (red data points) or a modified Hertz model that includes a linear term for the solid surface tension (blue data points). An indentation speed of 1 μm/s and a setpoint force of 10 nN were used for the measuerements. The x marks indicate independent experiments, the circles mean values and error bars SEM. **B)** Apparent Young’s moduli determined with the standard Hertz model as a function of indentation speed for elastomers of varying ζ. Data represent the average and error bars the SD from 75 measurements at different locations of the elastomers (one of two independent experiments shown). Data were fitted with a power law equation, with the fitted parameters given in the table. **C)** Apparent Young’s moduli determined with the standard Hertz model as a function of indentation depth for elastomers of varying ζ. The inset shows the data as a function of setpoint force. Data represent the average and error bars the SD from 75 measurements (one of two independent experiments shown). **D)** A typical F-d curve derived from indentation of an elastomer with ζ=0.7 along with best fits for the standard Hertz model and a modified Hertz model that includes a linear surface tension term. **E)** Solid surface tension of silicone elastomers derived from the modified Hertz model. The x marks indicate independent experiments and the column the average value.

*E* values from the indentation experiments were higher compared to those obtained with rheology assuming a Poisson ration of 0.495 (44), with differences being more pronounced for the softer elastomers (**Supplementary Fig. S2D)**. A close look of the Hertz model fits of the F-d curves for elastomers with ζ=0.7 showed important deviation from the experimental data (**Fig. 2D**). We reasoned that this could be due to an important contribution of solid surface tension (*G*) of soft silicone elastomers, as previously reported (44). When analyzed using a modified Hertz model that incorporates a linear surface tension term (see details in Methods section), the fit quality for the F-d curves was greatly improved, especially for elastomers with the lower ζ values (**Fig. 2D**), consistent with the expected larger contribution of surface tension for the softer substrates. The calculated values for *G* range between 3 and 10 mN/m, in agreement with previous reports (44) (**Fig. 2E**). In some cases, the variation in *E* was large when *G* was left as a fitted parameter; we therefore set as fixed value for *G* the mean calculated value, to obtain less scattering (**Supplementary Fig. S2E**). Using the modified Hertz model, the values of *E* calculated from 3 different batches were lower and closer to those obtained by rheology (**Fig. 2A**). Taken together, our results suggest that surface tension contributes significantly to indentation forces of soft elastomers and raise the question of how the apparent higher surface stiffness affects cell behavior in combination with bulk viscoelasticity.

### Fibronectin (FN) functionalization of elastomers

The hydrophobic character of elastomers allowed coating through simple adsorption of FN from solution. Remaining area between FN was blocked with albumin. The amount of adsorbed FN reached a plateau above coating concentrations of approx. 5 μg/ml, as measured using a modified ELISA assay (**Fig. 3A**). The number of adsorbed FN molecules was similar between elastomers of differing ζ ratio (**Fig. 3B**), an important prerequisite to attribute changes in cell behavior to substrate mechanical properties. Interestingly, accessibility to the C-terminal heparin-II binding (Hep2) and central cell binding (CBD) domains was higher for FN immobilized on elastomers compared to TCPS, indicating higher binding affinity and/or altered conformation, favoring interactions with cells (**Fig. 3B**). At the resolution of optical microscopy, fluorescently-labeled FN (FFN) formed a homogeneous coating, with no apparent assembly of fibrils or aggregation (**Supplementary Fig. S3A**). The ability of cells to remodel FN depends on the adhesion strength of FN to its underlying substrate, as well as the physical state of FN (48, 49). In order to alter the latter, FN-coated elastomers were treated with the fixation agent paraformaldehyde (PFA). PFA treatment did not alter the amount of adsorbed FN (**Fig. 3C**), as previously shown (49). Taken together, these data show the feasibility of preparing FN-coated elastomers of varying mechanical properties, but similar biochemical properties.

**Figure 3.**
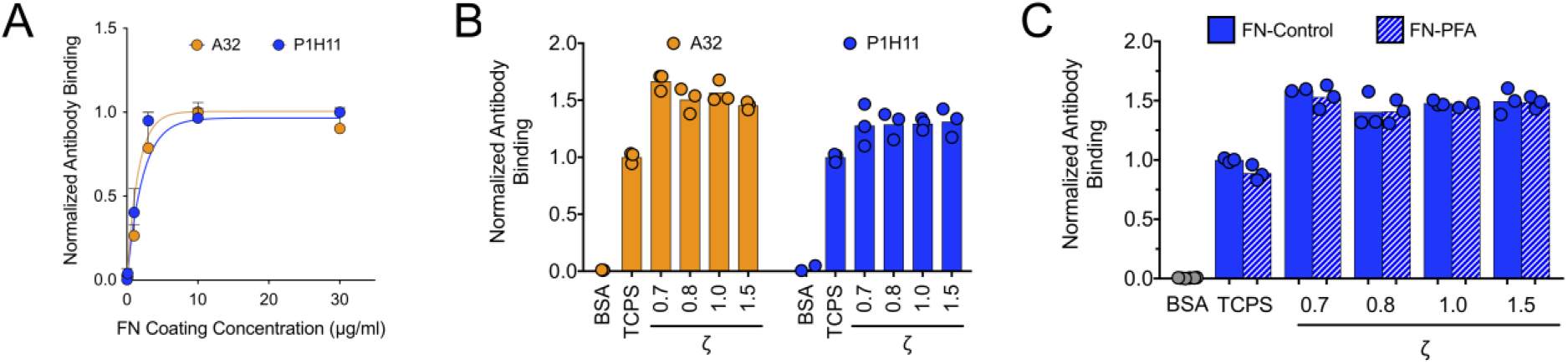
Fibronectin coats elastomers with similar efficiency. **A)** Coating efficiency of adsorbed FN on untreated silicone elastomers (ζ=1.0) from FN solutions of different concentration. FN was detected by a modified ELISA assay using two different monoclonal anti-FN antibodies: clone P1H11 recognizes the central cell binding domain and clone A32 the heparin II binding domain. Data from 2 independent experiments are presented: 3 replicates were measured in each experiment. Lines represent fits of the equation *y* = *A*(1 – *e*^−*kx*^). **B**) Coating efficiency from a 10 μg/ml FN solution on untreated silicone elastomers of varying ζ, TCPS and BSA-coated TCPS. FN adsorption to the substrate was similar on elastomers of different mechanical properties and higher compared to TCPS. Data from 1 (of 2) independent experiment are presented: each data point represents one measurement and the column the mean value. **C**) Coating efficiency from a 10 μg/ml FN solution on FN-coated substrates, untreated (control) or treated with 4% PFA to cross-link FN. Antibody (clone P1H11) binding to immobilized FN was not affected by PFA treatment. Data from 1 (of 2) independent experiment are presented: each data point represents one measurement and the column the mean value.

### Fibroblast polarization, but not spreading, depend on elastomer stiffness

We next examined how primary human dermal fibroblasts (pHDF) adhered on FN-coated elastomers as a function of their mechanical properties (ζ ratio), in an effort to comprehend and specify which material properties of the substrate regulate cell response. Previous work has correlated the ability of cells to spread efficiently and assemble mature focal adhesions (FAs) to substrate stiffness. However, this conclusion was drawn primarily from studies on highly elastic hydrogels (29, 50-52), which do not exhibit the frequency dependence in elastic moduli (**Fig. 1**) and apparently different surface, compared to bulk, mechanical properties (**Fig. 2**), like the elastomers described here. Indeed, studies on elastomer have reported mixed results concerning the effect of substrate mechanics on cell adhesion (9, 10, 12, 53); hence, the need to clarify how cells respond to the well-characterized elastomers reported here.

pHDF adhered with high efficiency on FN-coated elastomers, independent of their mechanical properties, and similar to traditional FN-coated TCPS (**Fig. 4A**). Interestingly, fibroblasts spread to the same extent on soft and stiff elastomers (**Fig. 4B-D**), while their aspect ratio, which reflects cell polarization, increased with ζ (**Fig. 4B,E**). Indeed, high resolution imaging of the actin cytoskeleton confirmed that on the softer elastomers (ζ = 0.7, 0.8), fibroblasts remained round, with filamentous actin assembled primarily in transverse arcs and dorsal stress fibers, whereas on the stiffer elastomers (ζ = 1.0, 1.5), pronounced, oriented ventral stress fibers were visible (**Fig. 4C**). Similarly, the microtubule network was polarized only on the stiffer elastomers (**Fig. 4C**). Focal adhesion morphology was evaluated by immunofluorescence microscopy of paxillin (**Fig 4C**). Fibroblasts assembled large, elongated FAs on all elastomers, independent of ζ, and similar to those assembled on rigid, FN-coated glass (**Fig. 4F,G**).

**Figure 4.**
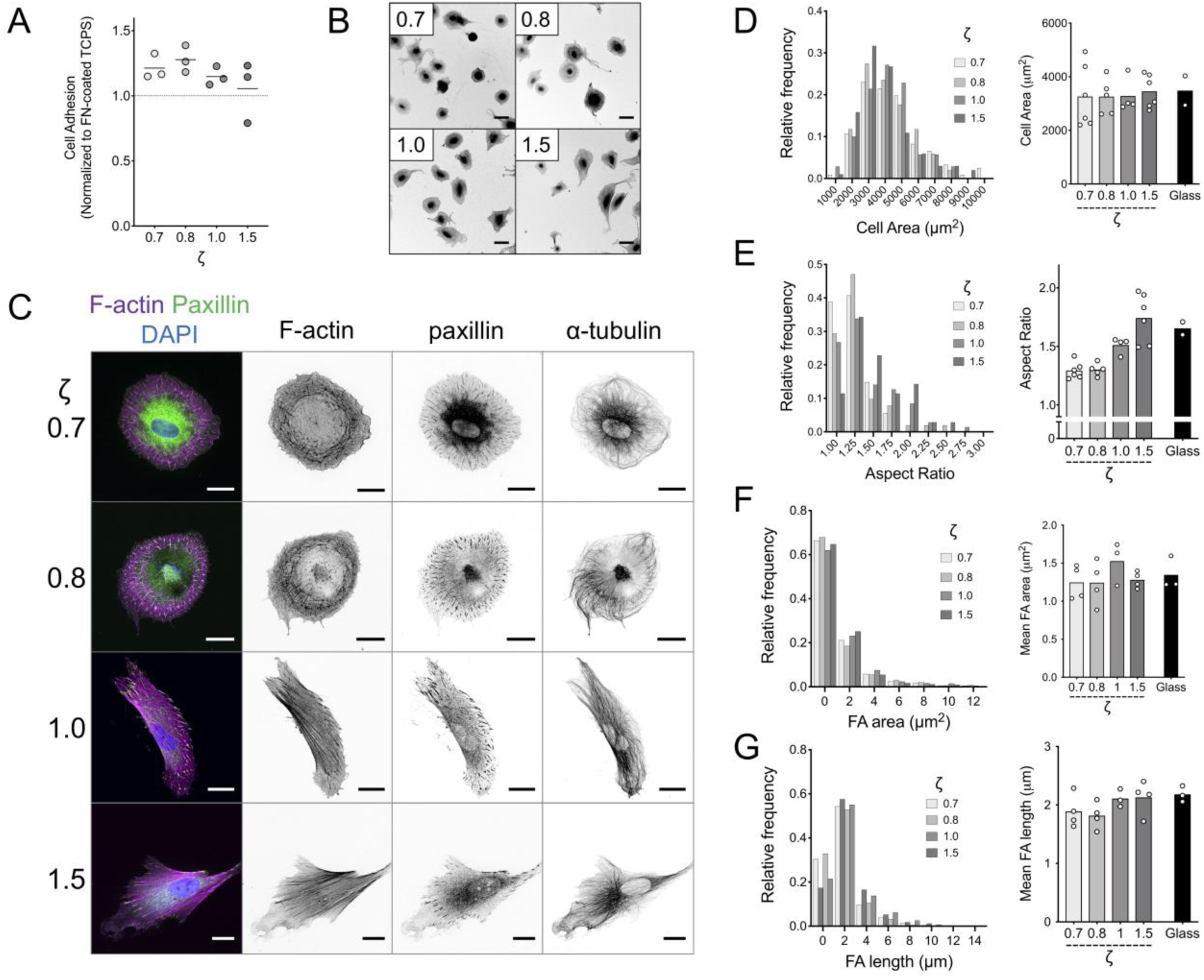
Elastomer mechanics regulates fibroblast polarization, but not spreading area or FA size. **A)** Adhesion of pHDF on FN-coated elastomers, normalized to their adhesion on FN-coated TCPS, as a function of ζ. **B)** Low magnification confocal microscopy images of fixed pHDF seeded on FN-coated elastomers for 4 hours and stained against F-actin using fluorescent Phalloidin to highlight overall cell morphology. Scale bars: 50 μm. **C)** High magnification confocal microscopy images of fixed and (immuno)stained pHDF seeded on FN-coated elastomer for 4 hours. Scale bars: 20 μm. **D-G)** Histograms showing the distribution of cell area (**D**), aspect ratio (**E**), FA area (**F**) and FA length (**G**) of pHDF seeded on FN-coated elastomer derived from one independent experiment as a function of ζ, as well as column plots showing the mean values from different independent experiments. FN-coated Glass was used as a control.

In sum, the viscoelastic properties of elastomers, in the studied range, did not affect the extent of FA and cell spread area, while elevated stiffness was required for fibroblast polarization. These findings suggest that either a different force threshold exists for different cellular processes or that cells employ different mechanisms to mechanosense and respond depending on the length scale and process (FA assembly *vs* polarization) involved.

### Fibroblasts do not remodel adsorbed FN on elastomers

In order for fibroblasts to sense the bulk substrate viscoelasticity, a sufficiently strong mechanical link between the adsorbed FN and the underlying elastomers must exist, so that FN is not simply removed from the surface. Fibroblasts can remodel physically-adsorbed FN on glass and other hydrophilic substrates (48, 54) raising the possibility that cell-generated forces could also result in FN unfolding and/or remodeling on elastomers. pHDF seeded on glass coated with fluorescently-labeled FN (FFN) extensively remodeled FN, as expected (**Supplementary Fig. S3B**): FFN was assembled into fibrils and some of it was internalized by cells (**Supplementary Fig. S3B,C & Supplementary Movie 2)**. FFN remodeling was largely inhibited after treatment of FFN with PFA (**Supplementary Fig. S3B**).

On elastomers, there was surprisingly no visual evidence of FFN fibrillogenesis or uptake as monitored by live-cell, epifluorescence microscopy. On the softer FFN-coated elastomers (ζ = 0.7) a striking accumulation of FFN under spreading cells, with darker areas around the cell periphery was observed (**Fig. 5A & Supplementary Movie 3**). On the stiffer elastomers, this effect was attenuated (ζ=1.0) or not observed (ζ=1.5) (**Fig. 5A & Supplementary Movies 4,5**). The increase in fluorescence intensity under cells over time was quantified and verified microscopy observations (**Fig. 5B**). Live-cell confocal microscopy of FFN at the surface plane of the softer elastomers (ζ=0.7) sometimes displayed dark areas under the cell body (**Supplementary Movie 6**). Confocal z-stacks showed that FFN was at a lower imaging plane, indicating that fibroblasts were markedly deforming elastomers, forming a crater-like structure (**Fig. 5C**). On the stiffer elastomers (ζ=1.5), deformations of the homogeneous FFN coating were hardly visible (**Fig. 5C**). Remarkably, fibroblasts were unable to assemble FN fibrils using the adsorbed FFN (**Fig. 5A,C**), even though they were still able to exert the necessary tractions to assemble fibrils containing endogenous FN (**Supplementary Fig. S4**). The above results demonstrate that cell-generated forces did not lead to remodeling of adsorbed FN on elastomers; instead, these forces were transmitted through the adhesive coating and led to stiffness-dependent elastomer deformation.

**Figure 5.**
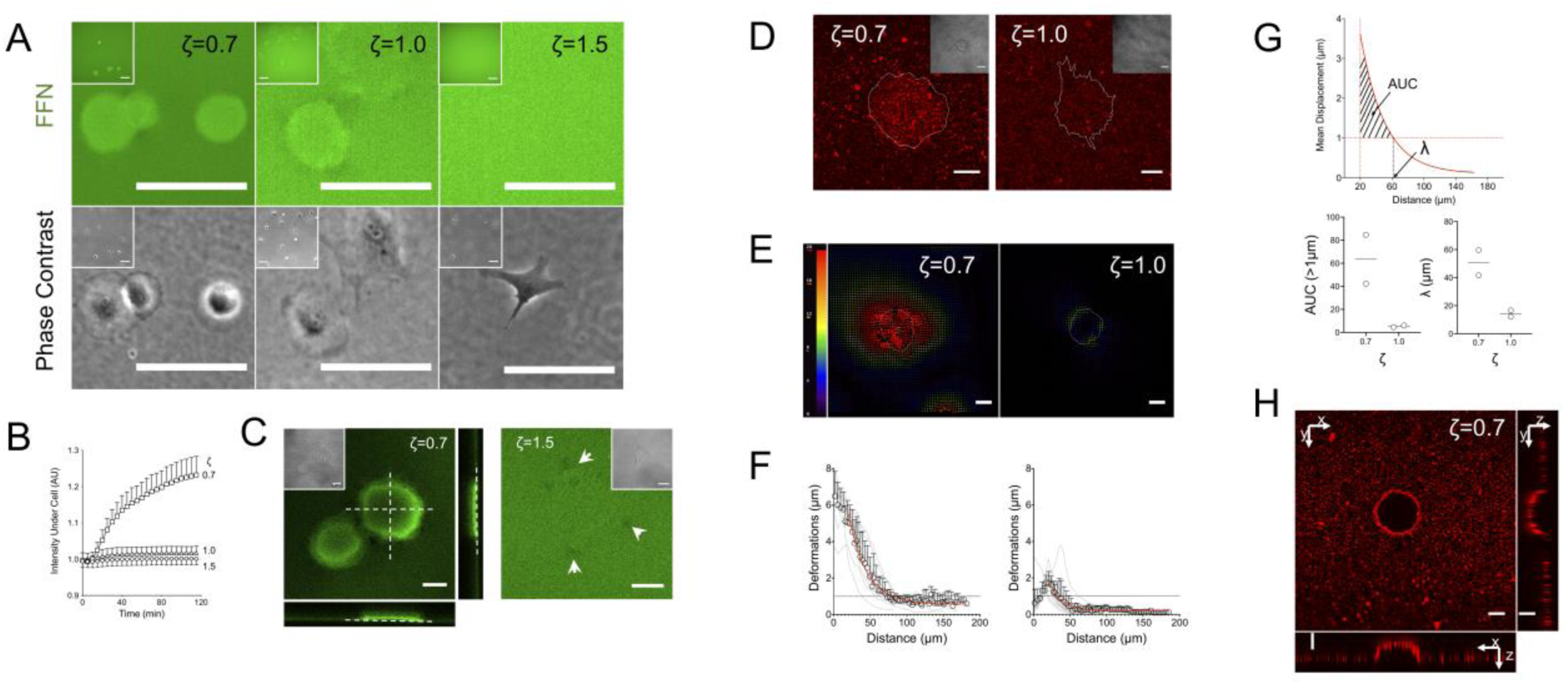
Fibroblast traction forces lead to substrate deformations but not fibronectin fibril assembly. **A)** Epifluorescence and phase contrast images of live pHDF seeded for 3 hours on elastomers coated with fluorescently-labeled FN (FFN). The intensity of FFN increased under the cell body on the softer (ζ=0.7) elastomers. **B)** Normalized FFN fluorescence intensity under the cell body normalized against background FFN intensity. Mean and SD from 2 independent experiments are presented (n=10). **C)** Confocal microscopy images of live pHDF seeded on FFN-coated elastomers. On the softer elastomers (ζ=0.7), surface out-of-plane deformations were observed. Dashed lines and arrows are visual aids. **D)** Confocal microscopy images of live pHDF seeded for 2 hours on elastomers with immobilized fluorescent beads and coated with FN. **E)** Particle image velocimetry (PIV) analysis from bead displacements on elastomers seeded with pHDF for 1 hour was used to produce substrate deformation fields. The magnitude of the color-coded vectors is given in pixels and the white line denotes the cell outline. **F)** Deformation as a function of distance from the cell center was calculated as detailed in the materials and methods section for each cell (grey lines) and averaged (circles with standard deviation). The data (mean ± SD, n=9 for ζ=0.7 and n=14 for ζ=1.0) were fitted using an exponential decay function (red line) starting from a distance of 20 μm, which corresponds to the typical cell radius. One of two independent experiments is presented. **G)** Quantification of two parameters reflecting total substrate deformation and the deformation boundary from the cell edge based on the fitted curves from (**F**). A threshold value of 1 μm for the deformation vectors was selected to calculate the distance from the cell edge at which deformations drop below the threshold and the area under the fitted curve (AUC) correlating with total substrate deformation. Both the boundary distance and AUC were lower for the stiffer elastomers. Each data point corresponds to an independent experiment and the line the mean. **H)** Confocal microscopy image of beads immobilized on a soft elastomer (ζ=0.7), which is being deformed by a living cell. Orthogonal views are presented to better visualize the crater-like structure formed. Scale bars 50 μm for **(A)** and 20 μm for **(C,D,E,H)**.

### Large-scale deformations of soft elastomers

To confirm that FN accumulation was due to elastomer surface deformation and to examine an alternative way to visualize this process, we immobilized fluorescent beads as fiducial markers on the elastomer surface, which was then coated with FN (55). Beads accumulated rapidly under fibroblasts seeded on the softer elastomers (ζ=0.7), similar to what was observed for labeled FN (**Fig. 5D & Supplementary Movie 7**). Again, there was no evidence for bead removal/internalization from the substrate in any of the data sets analyzed. Time-lapse imaging during cell spreading showed bead displacements over tens of micrometers towards cells on soft elastomers (ζ=0.7), indicating long-range deformations (**Supplementary Movie 8**). On the stiffer elastomers (ζ=1.0), beads were pulled towards cells as a result of traction forces, but exhibited much lower magnitude of deformations (**Supplementary Movie 9)**. We applied a PIV algorithm to visualize and quantify substrate deformations (**Supplementary Movies 10,11**); the displacement fields calculated from images before and 60 minutes after cell seeding were used to determine the radial deformation profile for cells on elastomers (**Fig. 5E,F**). Cell-induced deformations were much larger for the softer elastomers and extended further away from the cell body (**Fig. 5E,F)**. Interestingly, large deformations were also calculated under the cells for the softer elastomers, whereas they were maximal at the cell edge for the stiffer elastomers and decayed both towards the cell center and away from the cell. The decrease in substrate deformation was well-fitted with an exponential decay from the cell edge, assuming an average cell radius of 20 μm (**Fig. 5F)**. Defining a deformation field boundary at a threshold of 1 μm displacement, we measured a boundary distance of approx. 50 μm and 15 μm for the soft and stiff elastomers, respectively (**Fig. 5G)**. As a measure of total strain, we calculated the area under the fitted curve (AUC) from the cell edge till it intercepted the y=1 line (corresponding to 1 μm displacement), which confirmed the higher deformation on the softer elastomers (**Fig. 5G)**. In addition, during live-cell microscopy on the softer elastomers (ζ=0.7), out-of-plane deformation and the creation of a crater-like structure was often noted, similar to what was observed for fluorescent FN-coated elastomers (**Fig. 5H** & **Supplementary Movie 12)**. Fixation of cells for further microscopy analysis resulted in the disappearance, or attenuation of these structures, presumably due to traction force relaxation.

Overall, the above results demonstrated that cells generate large strains and deformations on the surface of soft elastomers. In combination with the presence of flow-like bead movement, this raised concerns for surface mobility of the coating and the presence of plastic deformations. Fluorescence recovery after photobleaching (FRAP) experiments of fluorescent fibronectin on the elastomer substrate showed absence of FFN recovery over hours, indicating that FN is not mobile on the elastomer surface and moves only after application of cell-generated forces (**Supplementary Fig. S5**).

We next examined whether cell-induced deformations were reversible upon cell death. Application of a 1% Triton-X aqueous solution resulted in rapid cell death; the recovery of FFN to its original homogeneous distribution prior to cell attachment was practically complete for cells on the stiffer elastomers, but only partial on the softer ones as evidenced by the inhomogeneous staining, indicating local plastic deformations of the coating (**Supplementary Fig. S6A**). Accordingly, bead accumulation under cells was still evident after Triton-X treatment on soft (ζ=0.7) elastomers, while beads recovered their initial positions on stiffer substrates (ζ=1.0) (**Supplementary Fig. S6B**). Occasionally, cell death occurred from inadvertent phototoxic effects, with the same result: partial recovery of beads to their original positions on the softer elastomers, thus excluding a specific effect of Triton-X (**Supplementary Fig. S6C**). The above results demonstrate that cells induced local plastic deformations on the surface of soft elastomers at the micrometer scale, presumably due to the high sol fraction of these materials. The apparent contradiction with the the absence of bulk viscoplasticity (**Fig. 1F**) can be explained when considering the length scales involved: the elastomer consists of connected (cross-linked) polymer chains, which provide the observed macroscopic elasticity, while soluble oligomers can flow between the polymer network and thus can irreversibly translocate upon stress application at the microscale.

### Reduction of cell contractility allows cell polarization on soft elastomers

The presence of robust focal adhesions and actin stress fibers, combined with the large induced deformations led us to hypothesize that fibroblasts were unable to polarize on the softer elastomers because the substrate could not resist the applied traction forces. Indeed, when force generation was reduced through mild, blebbistatin-induced myosin inhibition, fibroblasts polarized on the softer elastomers, as indicated by an increase of their aspect ratio (**Fig. 6A,B,D**), and the alignment of their actin cytoskeleton (**Fig. 6A,B**). Blebbistatin was used at concentrations that did not affect cell spread area (**Fig. 6C**), and did not impair assembly of mature FAs or stress fibers, which polarized along the main cell axis (**Fig. 6B**); nevertheless, blebbistatin-treated cells exhibited slightly smaller FAs (**Fig. 6E**). As expected, blebbistatin-treated cells induced smaller elastomer deformations, due to contractility inhibition. These counterintuitive findings showed that cells were able to polarize on soft elastomers when the forces transmitted to their substrate were lowered and suggest that substrate resistance to applied traction forces regulates cell polarization.

**Figure 6.**
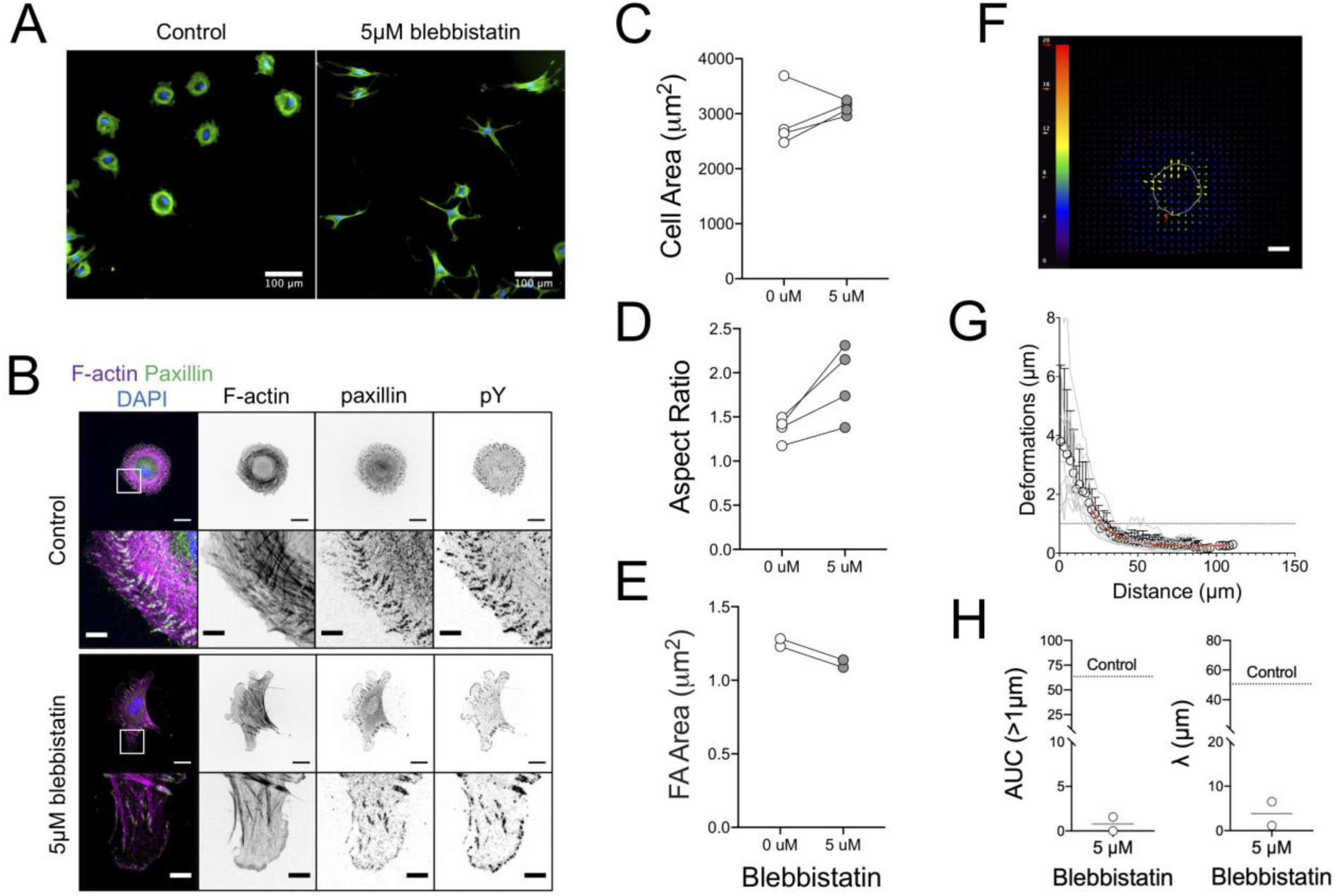
Inhibition of myosin II contractility results in reduced substrate deformation and fibroblast polarization. **A)** Epifluorescence microscopy images of Phalloidin-stained pHDF seeded for 3 hours on soft elastomers (ζ=0.7), treated with 5 μM blebbistatin or equivalent volume of carrier (DMSO). Nuclei are stained with DAPI (blue color). **B)** Confocal microscopy images of fixed and immunostained pHDF seeded on soft, FN-coated elastomers (ζ=0.7), treated with blebbistatin or DMSO. Scale bars: 20 μm/5 μm (details). Quantification of **C)** cell area, **D)** aspect ratio and **E)** FA area of pHDF cultured for 3 hours on soft elastomers (ζ=0.7), treated with 5 μM blebbistatin or equivalent volume of carrier (DMSO). Each data point represents the mean value of an independent experiment. **F)** Typical PIV analysis of bead displacements on an elastomer seeded with blebbistatin-treated pHDF for 1 hour. The magnitude of the color-coded vectors is given in pixels and is the same as for Figure 5E for comparison; the white line denotes the cell outline. Scale bar: 20 μm. **G)** Deformation as a function of distance from the cell center for blebbistatin-treated pHDF cells. The data (mean ± SD, n=8) were fitted using an exponential decay function (red line) starting from a distance of 20 μm, which corresponds to the typical cell radius. One of two independent experiments is presented. **G)** Quantification of the boundary distance and AUC for the situation of blebbistatin-treated cells. Comparison with the mean value calculated for control cells (dashed line; data from figure 5G) shows how contractility inhibition lowers substrate deformations.

## Discussion

Much controversy has troubled the field of mechanosensing following conflicting results from studies comparing cell responses on hydrogel versus silicone elastomer substrates of similar bulk stiffness (9, 12, 53). Our data suggest that one underlying reason for these discrepancies is the lack of thorough characterization of mechanical properties, which here revealed that soft elastomers exhibit a very important viscous component, giving rise to frequency-dependent stiffness, as well as a “stiff” surface layer, which we attribute to surface tension of the functionalized elastomers. Both of these factors are likely to contribute to the observed phenotypes. Since the stiffness sensed by cells can depend on the dynamics (frequency) of stress generation, it is likely that the material will appear soft or stiff depending on how fast cells exert traction forces on it. The presence of solid surface tension giving rise to a stiff surface layer is also expected to regulate cell behavior as evidenced by the ability of fibroblasts to spread efficiently and assemble FAs even on very compliant elastomers. This result resonates with some old - and recently revisited - studies, which showed that cells can be cultured even on liquid interfaces when these are stabilized with stiff protein films, despite the lack of an elastic character of the underlying oil (56, 57).

Our results argue against the direct comparison between findings obtained from cell mechanosensing studies on dissimilar materials based on single bulk stiffness values alone, and call for a thorough, comprehensive characterization of rubber materials, given the large number of developed silicone elastomer formulations (43, 58-60), including preformed, commercial substrates with nominal elasticities (e.g. www.softsubstrates.com). Notably, our findings demonstrated that different elastomer mechanical properties are derived depending on the technique used, supporting conclusions from previous studies (13, 58, 61), and highlighting the need for multiple dynamic measurements.

An often used and widely accepted measure of cell mechanosensitivity is the ability of cells to regulate the extent of spreading and assembly of integrin-based focal adhesions. Nevertheless, some findings argue that this measure is inadequate. *Firstly*, these two processes can be decoupled with cells spreading over large areas with small adhesions (27). *Secondly*, large traction forces are not necessary for assembly of mature FAs (15) and adhesion size does not always correlate with exerted tractions (62). In our study, fibroblasts exhibited similar spreading and FA assembly, independent of mechanical properties. Previously, studies have reported both an independence of cell area on substrate elasticity (9, 10), and a decrease in cell area on softer elastomers (53). Moreover, a decrease in FA area was noted for “softer” elastomers in one study (10). Important differences in the exact material formulation and corresponding mechanical properties can explain the above discrepancies: for example, in the aforementioned study (10), a different formulation (Sylgard 184) cross-linked at a very low ratio (1:75) of its components, with a nominal value of 5 kPa was considered the soft material, and the FA area was compared to a 2MPa substrate, which is >1 order of magnitude higher than the range studied here.

In contrast to spread area, important differences in cytoskeletal and cell polarization were observed as a function of elastomer viscoelasticity (**Fig. 4**). We argue that the inability of cells to break their original symmetry and randomly polarize on the softer elastomers was because these substrates could not resist the applied traction exerted on the material, through the tightly-coupled FN coating. Instead, the material flowed towards the cell and exhibited out-of-plane deformations. While the magnitude of traction forces was not directly measured, due to the challenges relating to the highly viscous component of the material and some surface viscoplasticity, the presence of stress fibers, large FAs and endogenous FN fibrillization all indicated that considerable forces were generated on the substrate, in contrast to what was observed on fluid, supported lipid bilayers (63, 64). Indeed, mild reduction of myosin II activity enabled cells to polarize even on the softest materials, which in this case resisted the attenuated applied forces. We thus propose that cell polarization on soft, deformable materials depends on the ability of the substrate to counterbalance applied traction forces, and thus provide sufficient friction to align actin stress fibers and form a force dipole at the cell scale along a predominant orientation (65). In other words, it is not the absolute mechanical properties of the substrate that determine polarization, but the force balance between the cell and the substrate.

We propose that the overall behavior of fibroblast on elastomers can be explained by considering two different processes of mechanosensing that differ substantially in length and time scales: 1) maturation of FAs at the subcellular scale through recruitment of integrins and associated adhesome proteins, and 2) cell polarization through cell-scale traction force generation. We suggest that the first process occurs through local changes brought about by rapid force loading on individual integrins and adaptor proteins, such that the substrate appears stiff, while the second process proceeds through probing of substrate mechanics by slower processes that involve larger force application.

Current models of substrate mechanosensing at the FA level assert that concurrent integrin engagement of the substrate and actin filaments through force-sensitive adaptor proteins underlies the fate of force transmission and adhesion cluster maturation (50, 66, 67). These “molecular clutch” models recently evolved to account also for a viscous element, revealing that viscosity indeed contributes to cell response depending on the timescales involved, and especially at lower substrate elasticities (64, 68). Integrins engaged simultaneously with the ECM and actin will be subjected to forces arising from actin retrograde flow in the order of 10-100 nm/s (50, 67, 69). Assuming a soft substrate with a linear spring constant of 1 pN/nm, this flow translates to loading rates on the integrins and adaptor proteins in the order of 10-100 pN/s while the “clutch” is engaged. At these high rates, and assuming an average force per integrin-actin linkage of a few pN (70, 71), e.g. 5 pN, integrins will probe the elastomer surface in less than a second, which will thus “appear” stiff based on the frequency dependence of measured elasticity (2-20 Hz, **Fig. 1**). Thus, forces could stabilize integrin-FN bonds, and lead to FA maturation; importantly, these forces may be myosin-II independent and instead stem from actin polymerization dynamics (52).

Clutch-based models rely on molecular interactions and mechanisms at the scale of FAs and thus cannot alone explain the process of polarization at the cellular scale. Several studies have suggested that mechanosensing emerges instead from larger-scale mechanisms that integrate information from cytoskeletal forces over the dimension of the cell (62, 72, 73). Cells pull periodically on their ligands using multiple FAs and thus related forces arising from multiple cytoskeleton-integrin-ECM linkages are substantially higher. Moreover, cells use a lower frequency range to probe their substrate at this length scale compared to that described for FAs above (74-76); at these lower frequencies (0.01-0.1 Hz), the elastomer would appear softer (**Fig. 1**) and would not resist the forces applied by cells, but instead flow as indeed was observed. Obviously, for this cell-scale mechanosensing to occur, initial assembly and maturation of FAs is a prerequisite. Therefore, our proposal is compatible with previously proposed local and global models of cell mechanosensing.

We recognize some potential complications of attributing the phenotype of cells on the ultrasoft elastomers solely on substrate mechanical properties. First, cells deformed the three dimensional morphology of the substrate at the cell scale, modifying microtopography, which is known to effect cell behavior on its own right (77). Second, the ligand density was altered as a result of substrate deformation, another effect that is expected to alter the level of intracellular signaling (78). Future work using micropatterned, adhesive islands, separated by non-adhesive regions will address these limitations, by inhibiting both processes.

In summary, the results of our study using well-characterized silicone elastomers suggest that cell mechanosensing is a multi-scale process. Hence, efforts should be devoted to bridge different scales in models, account for substrate viscosity and deformability, and provide a thorough dynamic characterization of substrates used in mechanosensing studies.

## Methods

### Reagents

A complete list of the commercially-available chemicals and antibodies used in this study are presented in Supplementary Tables S1, S2, respectively. Fluorescent fibronectin was prepared using bovine plasma fibronectin (Life Technologies, Cat. No. 33010018) and an AlexaFluor® 488 labeling kit (Thermo Fisher, Cat. No. A10235) according to the instructions provided.

### Elastomer Substrates

The silicone-based, 2-component elastomer formulation CY52-276 from Dow-Corning was used to prepared elastomers of varying mechanical properties by changing the weight ratio, ζ (A/B), of its two components. Component A (Base) and B (Catalyst) were mixed by vigorous magnetic stirring in glass vials for 5 minutes, degassed in vacuum for 2 minutes and the mixture was then applied, or spin-coated, onto different underlying substrates. The elastomer was cured for 3 hours at 65-70 °C, and then stored at room temperature until use. Elastomer substrates were used within 6 months.

Fibronectin (FN) was coated on elastomers through overnight incubation with freshly-prepared PBS solutions at 4°C. In order to cross-link FN, FN-coated substrates were treated with 4% PFA in PBS for 15 minutes at room temperature. Substrates were washed three times with PBS and incubated with 1% BSA in PBS for 30 minutes at room temperature. Relative coating efficiencies were calculated through a modified ELISA assay. Briefly, FN-coated substrates were incubated with 0.1 μg/mL of antibodies against FN for 1 hour at room temperature, followed by washing with PBS and incubation with secondary antibodies coupled to horseradish peroxidase (HRP) (0.16 μg/mL) for 1 hour at room temperature. Wells were washed with PBS and secondary antibodies were detected using a TMB substrate (3,3’,5,5’-tetramethylbenzidine; Sigma #T5569) and absorbance measurements at 630 nm.

Immobilization of fluorescent nanoparticles (200 nm red carboxylate-modified Fluospheres™) on the elastomer surface was adapted from a previous study(55) and was performed as following: elastomers were treated for 1 hour at room temperature with APTES in ethanol (10% v/v) and a drop of triethylamine. Elastomers were then washed once with ethanol, twice with water, and incubated with an aqueous suspension of 8.2×10^10^ fluorescent nanoparticles and 1 mg/ml EDC for 1 hour at room temperature. Elastomers were then washed with water and PBS before being incubated with FN as detailed above.

In order to determine the sol fraction of the elastomers, pre-weighed samples (*W*_*bef*_) formed on glass coverslips were incubated for 24 hours in n-hexane, washed with n-hexane, dried overnight and then for 1 hour under vacuum, before measuring their final weight (*W*_*gel*_). The weight of the sol fraction (*W*_*sol*_) was calculated as *Wsol* = *Wbef* – *Wgel* and the sol fraction as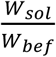.

### Rheometry

All rheometry measurements were performed on a Malvern Kinexus rheometer with parallel plate geometry and a temperature-controlled chamber. Temperature was controlled with an accuracy of 0.1°C. Elastomer components were mixed and degassed at room temperature as described above, and approximately 250 μl were applied on the bottom plate. The upper plate was then lowered until sample filled the gap between plates (gap size 500– 1000 mm). Initially a kinetic study was performed after heating the elastomer to 70°C and monitoring the storage modulus *G*^*′*^ (elastic modulus) and loss modulus *G*^*′′*^ (viscous modulus) in oscillatory mode for 3 hours. The temperature was then equilibrated at 25°C and a series of measurements in oscillatory mode were performed: 1) a frequency sweep (0.01-10 Hz), 2) a strain sweep (0.2-20%) and 3) creep measurements under different applied shear stress. All experiments were performed for at least three different batches.

### Atomic Force Microscopy

Silicone elastomers were characterized by indentation measurements using a Nano-Wizard III AFM (JPK Instruments AG, Germany). Cantilevers with a spherical, borosilicate glass probe (sQube®) of 5 μm in diameter were used. Cantilever spring constants were determined using the thermal noise calibration method and ranged between 0.45 – 0.60 N/m. Young’s Moduli were derived by fitting force-distance curves with a Hertz Model or a modified version of that model using a processing script written in Python (source code available upon demand) as described below:

Force distance curves were derived correcting the Z-piezo positions with the deflection of the curves. Background force were subtracted fitting a straight line to the end part of the curve. Contact point was first estimated taking the standard deviation (σ) of the end 10% of the curve, and estimating where the curve was first lower then –σ.

The Hertz-model of a spherical indenter: 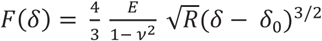 where d > 0 and zero everywhere else, was fitted to all values below a user specified maximal force using a non-linear least-squares method (least sq. method from SciPy, based on the Levenberg-Marquardt algorithm) to optimize 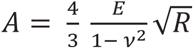 and d_0_. Where E stands for Young’s modulus, ν for the Poisson’s ratio of the sample, R is the radius of the indenting sphere, δ the indentation depth, and δ_0_ is its zero position. In the case considering the surface tension of the sample, the equation was modified to: 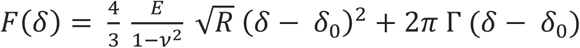. Here it was possible to fit for *B* = 2 π Γ as well or specify it as an a priori known constant.

### Cell Culture

Primary human dermal fibroblasts (pHDF) were purchased from ATCC (Cat # PCS-201-010) and cultured as sub-confluent monolayers at 37°C and 5% CO_2_, in Dulbecco’s modified eagle’s medium (DMEM; Life Technologies; Prod. # 10938), supplemented with 10% fetal bovine serum (FBS) and 1% penicillin/streptomycin (P/S). pHDF cultures were used until passage 15 and checked monthly for the absence of mycoplasma.

### Cell Adhesion Assays

The efficiency of cell adhesion was evaluated 30 minutes after seeding on coated substrates inside a 96-well plate. Trypsinized cells were kept in suspension under ice for 10 minutes before addition to wells (100 μl of 1×10^5^ cells/ml corresponding to 3.1×10^4^ cells/cm^2^). After 30 minutes, wells were washed twice with ice-cold PBS, the solution was aspirated and plates were placed at −80°C overnight. Relative cell numbers were quantified using the Cyquant cell proliferation assay kit (Thermo-Fischer). Of note, the sample solution above the substrates was moved to a new 96-well plate for fluorescence measurements, in order to avoid interference from the elastomer substrate.

### Microscopy

Immunofluorescence microscopy was performed on cells fixed with 4% PFA in PBS for 15 minutes at room temperature. Membranes were permeabilized by incubating with Triton X-100 (0.1%) for 5 minutes, followed by blocking with 1% BSA in PBS. Primary antibodies (diluted 1:100 in 1% BSA) were incubated for 1 hour at room temperature or overnight at 4°C. Cells were washed and incubated with secondary Alexa Fluor®-labeled antibodies (Life Technologies; diluted 1:150 in 1% BSA) for 1 hour at room temperature.

DAPI and TRITC-phalloidin were used to stain nuclei and filamentous actin (F-actin). Images were acquired on a Zeiss LSM 880 laser scanning confocal microscope (LSCM) using a 63x/1.4 NA oil-immersion objective (Zeiss) or a 20x/0.8 NA objective (Zeiss).

Live-cell, time-lapse microscopy on FFN-coated substrates or substrates with immobilized fluorescent beads was performed on the Zeiss LSM 880 confocal microscope or the Leica DMi8 microscope. For live-cell imaging, CO_2_-independent medium (Thermo Fisher Cat# 18045054) supplemented with 10% FBS and 1% P/S was used.

Fluorescence recovery after photobleaching (FRAP) analysis was performed using FFN-coated elastomers in PBS solutions on a Leica DMi8 microscope equipped with a 488 nm laser source and a 40x, NA=0.60 objective. A predefined spot of approx. 100 μm^2^ was bleached (within < 1 second) and the recovery was monitored over a period of 2 hours at one minute intervals or 10 hours at 5 minute intervals. Microscopy images were analyzed using ImageJ: the mean fluorescence intensity of the bleached region was measured, corrected for photobleaching due to imaging and normalized.

### Elastomer Deformation (TFM protocol)

In order to measure cell-induced substrate deformations, elastomers with immobilized fluorescent nanoparticles were employed. Substrates were first equilibrated at 37°C in the environmental stage of a Zeiss LSM 880 LSCM, in presence of CO_2_-independent medium. A small volume of pHDF were subsequently added to obtain a cell density of 5×10^3^ cells/cm^2^ and imaging was immediately set off. Hence, the first image was the reference state of the elastomer. Images were acquired using a water-immersion 40x objective (Zeiss, NA=1.20) at different time points after cell addition. In order to map substrate deformations, stacks of images before and after cell addition were first aligned using the StackReg plugin of ImageJ (http://bigwww.epfl.ch/thevenaz/stackreg/), followed by particle image velocimetry analysis performed using the PIV plugin (https://sites.google.com/site/qingzongtseng/piv). Vector maps were further analyzed as following:: The map data was imported into a python script (available upon demand), and converted to an image of strain field. This image was converted to a binary using threshold of 20% of the maximum displacement and individual patches were identified. This was necessary to handle cases where more than one cells were present on the image.

The center of each patch was estimated using a weighted mean, and the points converted to polar coordinates around this center. A radial displacement was calculated by binning the distances to a predefined set of radii (0-R_max_ pixels with single pixel steps) and calculating their averages and standard deviations. A linearized exponential fit was performed between the maximum and the first local minimum (if existed) behind it to estimate the decay of the profile.

### Image analysis

Cell projected area and aspect ratio, defined as the ratio of the major to the minor axis of a fitted ellipse, were determined through image analysis of single phalloidin-stained cells using the ‘Cell Outliner’ plugin of ImageJ. Focal adhesion area was determined using a custom-written macro in ImageJ, as previously described(51). An area threshold of 0.4 μm^2^ was set to exclude small focal complexes and noise.

## Author Contributions

D.M. designed the study, performed experiments, analyzed data, supervised the project and wrote the manuscript. T.H. analyzed experimental data and commented on the drafted manuscript. L.H. performed immunofluorescence microscopy and fibronectin ELISA assays. J.P.S. supervised, secured funding for the project, and commented on the drafted manuscript.

## Competing Interests

The authors declare no competing interests.

